# Probabilistic Annotations of Protein Sequences for Intrinsically Disordered Features

**DOI:** 10.1101/2024.12.18.629275

**Authors:** Nawar Malhis

## Abstract

This paper introduces a novel platform for IDR Probabilistic Annotation (IPA). The IPA platform now encompasses tools for predicting ‘Linker’ regions and ‘nucleic’, ‘protein’, and ‘all’ (protein or nucleic) IDR binding sites within protein amino acid sequences. Despite its simplicity and computational efficiency, results demonstrate that IPA performs competitively with leading tools in predicting ‘protein’ and ‘all’ IDR binding sites while considerably outperforming all tools in identifying Linker regions and nucleic binding sites. An important contribution of this work is the introduction of a new output paradigm for computational feature predictions. Traditional tools typically express predictions as scores, with higher values indicating greater probabilities. However, these scores lack true probabilistic meaning and interpretability, even derived from logistic regression models. This limitation arises primarily because training data priors differ from broader populations’ unknown priors. This paper proposes applying a reverse Bayes Rule to logistic regression outputs, effectively normalizing for the priors in the training data. This adjustment produces scores representing actual probabilities, assuming 50% priors in the general population. Such scores are interpretable in isolation and enable comparability and integration across different tools, marking a significant step toward standardization in feature prediction methodologies.

**Availability:** orca.msl.ubc.ca/nmshare/ipa.tar.gz

## 1. Introduction

Intrinsically disordered protein regions (IDRs) are segments of proteins that lack a fixed or stable three-dimensional structure under physiological conditions [1][2][3]. They are rich in polar and charged amino acids (e.g., Serine, Glutamine, Lysine) and deficient in hydrophobic residues that drive folding in structured proteins [4]. IDRs are prevalent in eukaryotic proteomes, with an estimated 20% of eukaryotic protein residues being disordered; many are associated with diverse biological functions [5].

IDR interactions are characterized by high specificity and low affinity, i.e., reversible [6], making them suitable for signaling pathways and regulation processes, including allosteric regulation [7]. Upon binding to folded protein domains, IDRs adopt conformations complementary to their binding partners; they often fold upon binding, a phenomenon known as coupled folding and binding [8][9]. Interactions between IDRs and other IDR partners display diversity in conformational outcomes, broadly categorized into two types: 1) fuzzy interactions, in which IDRs retain a significant degree of conformational flexibility in the bound state [10]. 2) folding upon binding, where IDRs undergo coupled folding and binding, transitioning from their disordered state to a well-defined structure [11]. This flexibility allows IDRs to recruit various binding partners, acting as hubs in protein interaction networks [12]. IDRs are often regulated by post-translational modification (PTM), such as phosphorylation, acetylation, and ubiquitination [13].

Annotating protein sequences with IDR interaction sites and other features is essential for understanding their roles in biological systems and their contribution to biological processes. Manually curated annotations sourced from experimental results are considered the highest-quality annotations. However, like other protein annotations, such annotations are limited and only cover a tiny proportion of the overall IDR features. Note that the percentage of the manually curated SwissProt sequences in the latest release of the UniProtKB database (2024_06) is 0.23% [14]. While SwissProt sequences are selected among those enriched in molecular functions, these sequences are not fully annotated. Thus, we can reasonably assume that the percentage of manually curated molecular function sites is comparable, i.e., ∼1%. Consequently, understanding unannotated protein sequences relies heavily on high-throughput computational tools [15 … 22].

Various computational tools have been developed for annotating protein sequences within intrinsically disordered regions [23 ‥ 50]. Several attempts have been made to understand and assess the prediction quality of these tools [51 ‥ 57], most notably the community-based Critical Assessment of protein Intrinsic Disorder prediction (CAID)[51][56]. CAID utilizes newly annotated sequences from the DisProt database [58] to evaluate the predictions of IDR and IDR-binding sites. CAID test datasets are fair as they only include new manually curated annotations released after the training of the evaluated predictors. My only concern with respect to the CAID test datasets is that they do not account for the known false negative sites by limiting their annotations to newly unreleased ones. For example, if we extend the annotations of the 3,022 DisProt (2024_06) sequences by the annotations of the MobiDB-LIP [59] database, the number of protein-binding residues increases by more than a third from 68,925 to 82,394.

Furthermore, DisProt annotations used are limited to two states, True/False, e.g., protein binding vs. non-binding. Including a third ‘unknown’ state to represent potentially True residues can help reduce false negatives. Marking low-quality annotations of protein-binding IDR from the MobiDB-LIP database and ‘ambiguous’ annotations of the DisProt, as described in **Section 2.2**, converts the annotation of a total of 237,656 residues from non-protein-binding to ‘unknown’, i.e., decreases the number of the ‘negative’ state residues by ∼14.5%.

Apart from the annotation, the non-standardized outputs of prediction tools can lead to a potential misinterpretation. Generally, most tools provide two output forms: an analog score and a binary prediction obtained by thresholding the analog score. The analog score is usually called a propensity score, and the definition of ‘propensity’, if provided, can vary, as it can be defined as probability [26] and used interchangeably or more broadly as a numerical value such that residues with higher propensities are more likely to be the targeted feature residues [49]. **Figure 1** illustrates the lack of uniformity in the distribution of IDR-protein binding propensity scores generated by three widely used prediction tools against the sequences in the TEST_24_06 dataset (section 2.2). This lack of uniformity limits the interpretation of these predictions to the relative likelihood of binding to protein targets, as the absolute score values provide little information. In addition, these predicted scores are not directly comparable. Binary predictions aim to simplify interpretation by offering definitive values. However, the absence of a standardized thresholding mechanism leads to different tools predicting significantly varying percentages of the input as positive residues, further complicating the interpretations. To overcome these issues, the authors of the CAID portal server [60] provided alternative threshold values, enabling the user to choose to replace the predictor’s default threshold with optimized thresholds corresponding to a selection of metrics.

**Figure 1:**
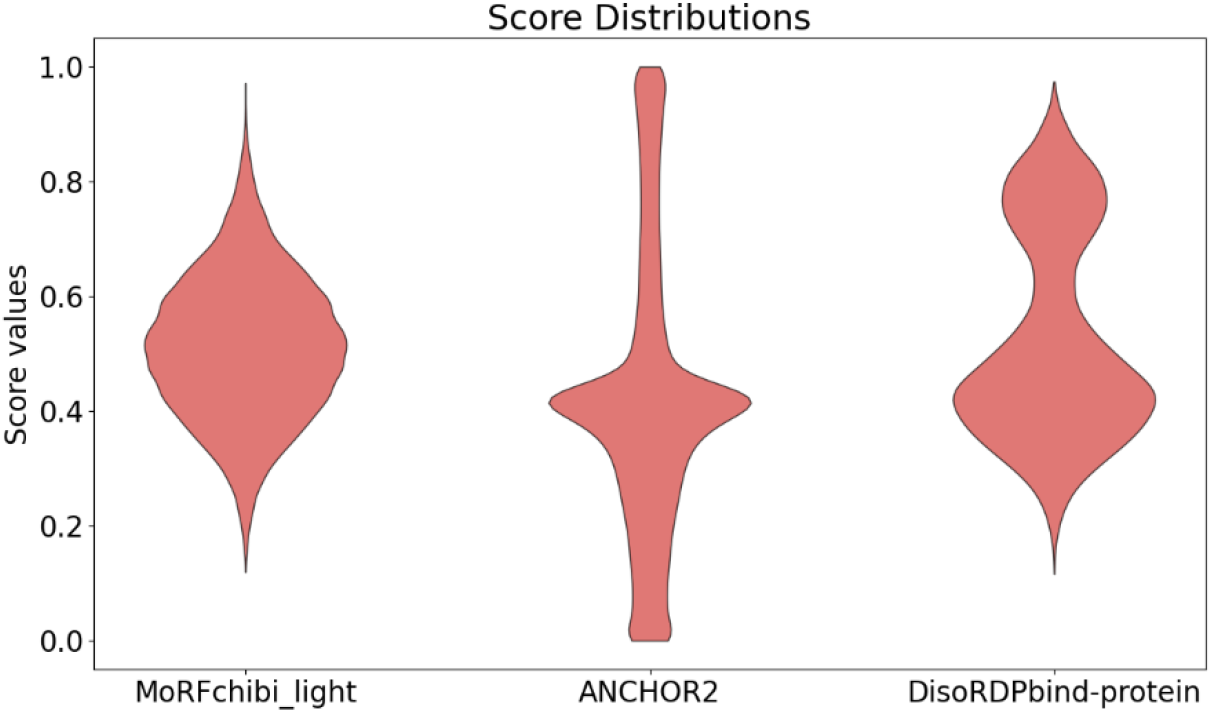
Violin plots for the propensity scores of MoRFchibi_light, ANCHOR2, and DisoRDPbind-protein against the **TEST-24_04** dataset, reflecting substantial inconsistencies of these scores.

In this paper, I present a platform for predicting annotations related to disordered protein regions from protein sequences. Because the prior for each category is unknown, i.e., the percentage of positive in the general population, I designed IPA predictions to generate, to the degree possible, probability values assuming 50% priors. Consequently, the scores of IPA predictions can be understood individually, not just as relative to other scores. And because IPA probabilities are simple and easily interpretable, IPA outputs do not include binary values. When binary predictions are needed, users can threshold IPA scores appropriately to their needs. At this point, IPA generates predictions for IDR protein-binding sites, IDR nucleic-binding sites, and IDR all-binding sites, where ‘all’ refers to nucleic or amino acids. In addition, I also created a second flavor of IPA that utilizes AlphaFold-2 predicted structures, thus called IPA-AF2, to predict protein-binding sites and Linker regions.

But before going into the details of IPA’s predictions, let’s first define IDR binding sites in general. For an IDR site to bind to any partner, the targeted partner must be available within the vicinity of the site, and other environmental conditions must be favorable, e.g., organism, subcellular location, temperature, atmospheric pressure, phosphorylation state, etc. However, tools are optimized solely based on the protein’s annotated amino acid sequences, and binding predictions are based solely on these sequences. Therefore, the best we can achieve is to predict sites with the potential of binding if everything else is available. Consequently, homologous sequences will have similar predictions, even though they might not exist within the correct species or the intended binding favorable condition. In other words, from the computational viewpoint, homologs to a protein binding site are protein binding sites regardless of their actual molecular function.

## 2. Results

Even though the scores generated by many tools are outputs of logistic regression models, these scores are not likely to represent actual probabilities because of the annotation incompleteness of the training datasets, and the composition of positives and negatives in these datasets are not expected to represent the unknown composition of the general population. Plus, and most importantly, the percentages of ‘positive’ in the training datasets, i.e., the priors, do not reflect those percentages in the general populations, which are also unknown. To reduce the effect of the annotation incompleteness and compositions to the degree possible, I assembled training and testing datasets by integrating the annotations of multiple databases into three-state annotation data. High-quality annotations represent the ‘Positive’ state, low-quality the ‘Unknown’ state, and the remaining residues are in the ‘Negative’ state. To resolve the unknown priors issue, I applied a reversed Bayes rule to the IPA outputs generated using logistic regression models to factor out the priors of the training data. Thus, IPA’s outputs are probabilities, assuming 50% priors.

### 2.1 Training datasets

I downloaded all 2,845 sequences of then (March 2024) the latest release, 2023_12, of the DisProt database and selected protein-binding (GO:0005515), nucleic acid, or just nucleic binding (GO:0003676, GO:0003723, and GO:0003677), and Linker regions (Intrinsically Disordered Proteins Ontology, IDPO:00502). I complemented the protein-binding annotations of the DisProt sequences with manually curated annotations from the IDEAL [61] and the DIBS [62] databases. I assembled a dataset with three-state annotations: manually curated annotations are in the ‘positive’ class, DisProt Ambiguous annotations are assigned to the ‘unknown’ class, and the remaining residues are in the ‘negative’ class. I used CH-HIT [63] to cluster sequences at 90% identity and only retained the cluster centers. The annotations of sequences at 100% identity are transferred to cluster centers. I used HAC [64] to mask, i.e., convert to ‘unknown’, conflicting annotations separately in each category. For training cross-validation (N=4), I divided the sequences for each of the ‘linker’, ‘protein’, ‘nucleotides’, and ‘all’ (protein and nucleic binding) IDR-binding annotations at random into four groups, such that the number of binding/Linker residues and the number of binding-sites and linker-regions are about equal in each group. Finally, for each of the four groups, I doubled (for ‘linker’, ‘protein’, and ‘all’) and tripled (for nucleic) its sequences by adding random sequences from the remaining DisProt sequences.

### 2.2 Test dataset

For the test dataset, I include all descendants of each GO term for each annotation category to increase the data size. First, I included all of the 185 new DisProt release 2024_06 sequences that are not in the 2023_12 release, **Supplementary Table S1** (DSET1). To eliminate potential data contamination related to test sequences homologous to those used in training, I used CD-HIT to filter out test sequences with more than 30% identity to those used in training, **Supplementary Table S1** (DSET2). Then, I extend the annotations of these sequences from the MobiDB-LIP database. MobiDB integrates manually curated IDR-binding annotations from FuzDB [10], IDEAL [61], DIBS [62], MFIB [11], ELM [65], and DisProt databases, homologous to these annotations, as well as high-throughput annotations derived from PDB structures by Flipper [66]. In line with my earlier discussion, MobiDB considers the quality of homology annotations comparable to those curated and much higher than that of the derived. Thus, I included all the curated and homology annotations for DIBS, IDEAL, and MFIB for protein binding and masked out the derived annotations. I also masked out the curated and the homology for FuzDB and ELM as they include non-protein binding sites, **Supplementary Table S1** (DSET3). While CD-HIT excluded sequences with more than 30% identity to the training data, short regions homolog to the training sequences can still exist in the test data. Thus, I used HAM [64] to mask out homolog regions equal to or longer than ten residues with an identity to the training sequences higher than 80%. **Supplementary Table S1** (DSET4). The resulting test dataset is small and insufficient to provide quality evaluation for the five IPA predictors, especially the linker regions and the nucleotide-binding sites. Thus, I also included all new 2024_06 nucleic-binding, protein-binding, and linker regions annotations to sequences in the 2023_12 release after masking out the remaining residues in these sequences; see an example in **Figure 2**.

**Figure 2:**
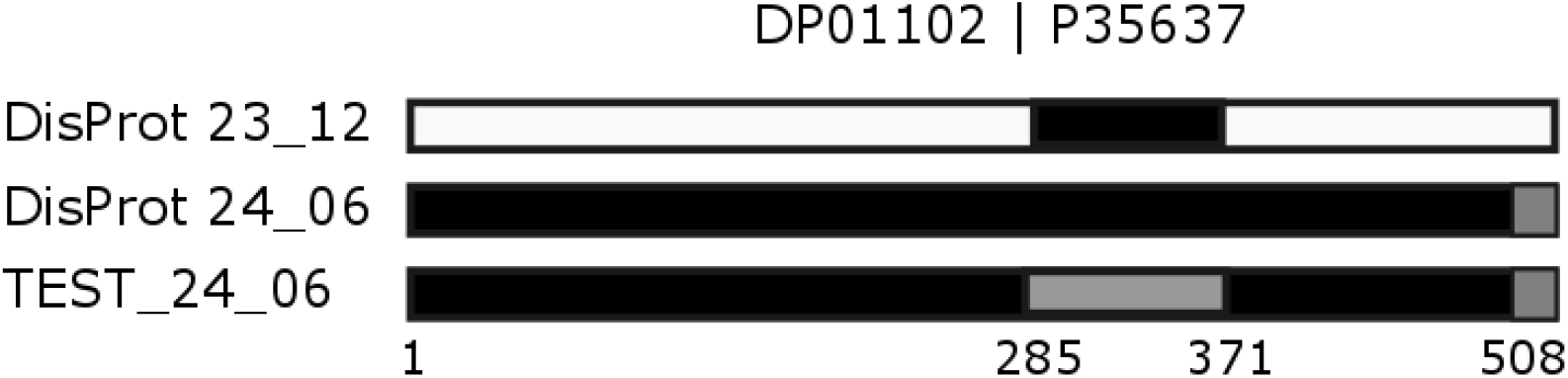
Annotation of nucleic-binding IDR residues in the DisProt DP01102 (UniProt P35637). DP01102 residues 285 to 370 were annotated as a nucleic-binding IDR site in the DisProt release 23_12 (black). DP01102 annotation is revised in the 24_06 release to include all 526 residues. However, residues from 508 to 526 are not confirmed to be disordered under physiological conditions. Thus, in the 24_06 release, residues from 1 to 507 are IDR-nucleic-binding (black), and 508 to 526 are ‘unknown’ (gray). TEST-24_06 nucleic-binding residues for DP01102 are those in the 24_06 release and not in the 23_12 release. Thus, nucleotide-binding residues are from 1 to 284 and 371 to 507; the remaining are ‘unknown’ (gray). ‘Black’ represents nucleic binding, ‘gray’ represents unknown, and ‘white’ represents non-nucleic binding.

The resulting TEST_24_06 dataset includes 184 sequences, 42 annotated with protein binding sites, 11 with nucleic binding sites, 48 with all binding sites, and 14 with linker regions—**supplementary Table S1** (TEST_24_06). IPA-AF2 predictors require structures generated by AlphaFold-2 from the AlphaFold database; thus, they can only process a subset of the TEST-24_06 sequences, TEST-24_06-AF2—**table 1**.

**Table 1.**
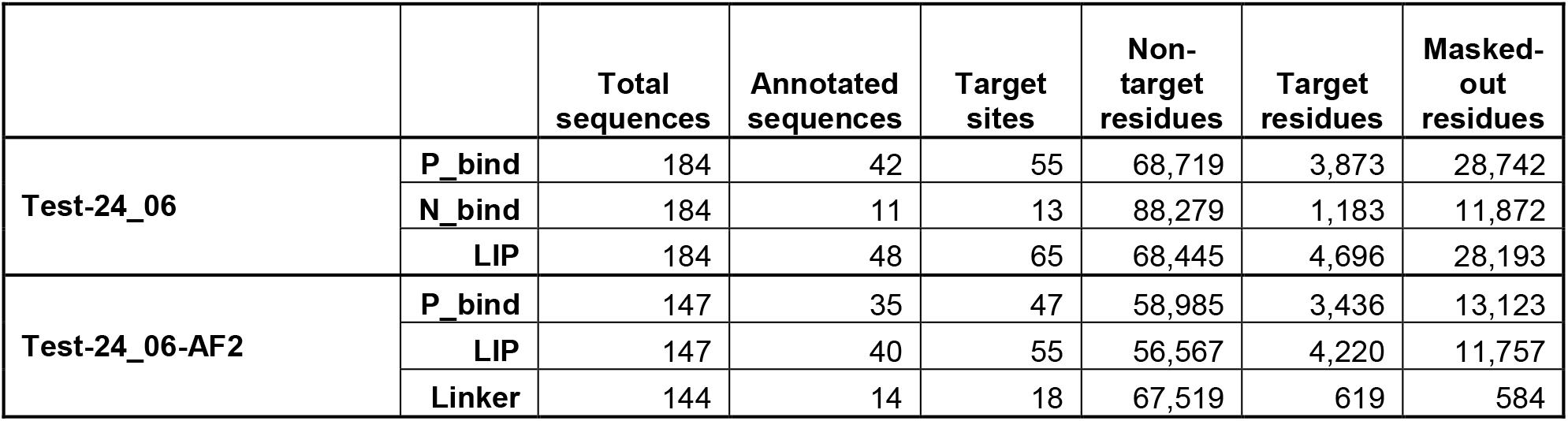
The **Test-24_06** dataset. **Test-24_06-AF2** is a subset of the TEST-24_06 sequences with structures generated by AlphaFold-2 in the AlphaFold database.

### 2.3 Comparison with existing predictors

I contrasted the predictions of each one of the IPA predictors with the leading relevant tools available against the TEST-24_06 dataset. **Figure 3** and **Supplementary Figure S1** show the ROC curves for the IPA and IPA-AF2 predictions compared to other tools. **Table 2** and **Supplementary Table S2** show that IPA and IPA-AF2 predictors outperformed or matched others in AUC values.

**Table 2.**
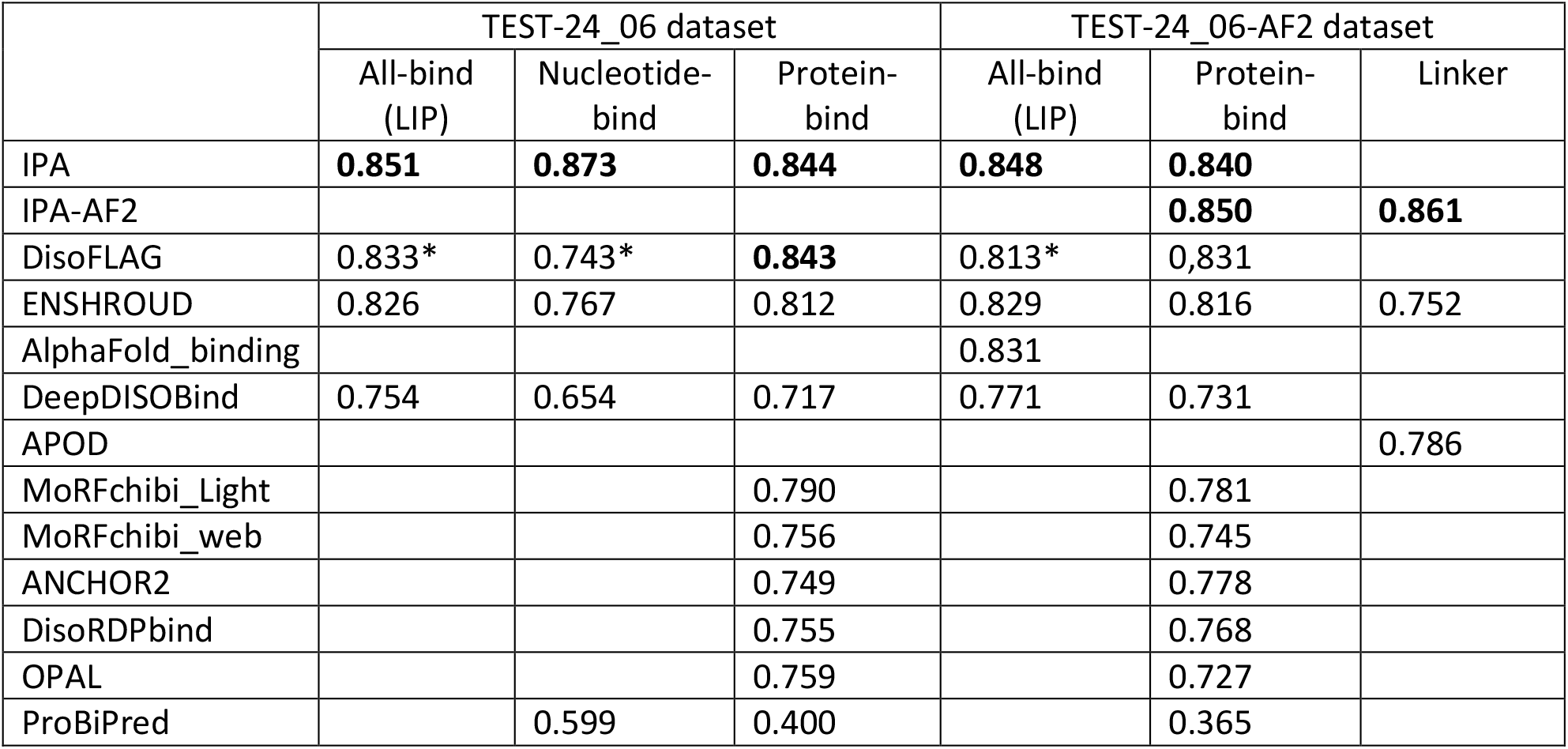
The AUC values for the IPA predictions compared to other tools. The highest AUC value and those within one AUC point are in bold for each column. (*) The DisoFLAG predictor does not generate predictions for nucleic-binding but generates predictions for RND-binding (RB) and DNA-binding (DB); thus, I averaged the scores of RB and DB as DisoFLAG-nucleic. It also does not generate predictions for all-binding; thus, I averaged the scores of DisoFLAG-nucleic and DisoFLAG-PB as DisoFLAG-all.

|  | TEST-24_06 dataset |  |  | TEST-24_06-AF2 dataset |  |  |
| --- | --- | --- | --- | --- | --- | --- |
|  | All-bind (LIP) | Nucleotide-bind | Protein-bind | All-bind (LIP) | Protein-bind | Linker |
| IPA | **0.851** | **0.873** | **0.844** | **0.848** | **0.840** |  |
| IPA-AF2 |  |  |  |  | **0.850** | **0.861** |
| DisoFLAG | 0.833* | 0.743* | **0.843** | 0.813* | 0.831 |  |
| ENSHROUD | 0.826 | 0.767 | 0.812 | 0.829 | 0.816 | 0.752 |
| AlphaFold_binding |  |  |  | 0.831 |  |  |
| DeepDISOBind | 0.754 | 0.654 | 0.717 | 0.771 | 0.731 |  |
| APOD |  |  |  |  |  | 0.786 |
| MoRFchibi_Light |  |  | 0.790 |  | 0.781 |  |
| MoRFchibi_web |  |  | 0.756 |  | 0.745 |  |
| ANCHOR2 |  |  | 0.749 |  | 0.778 |  |
| DisoRDPbind |  |  | 0.755 |  | 0.768 |  |
| OPAL |  |  | 0.759 |  | 0.727 |  |
| ProBiPred |  | 0.599 | 0.400 |  | 0.365 |  |

**Figure 3:**
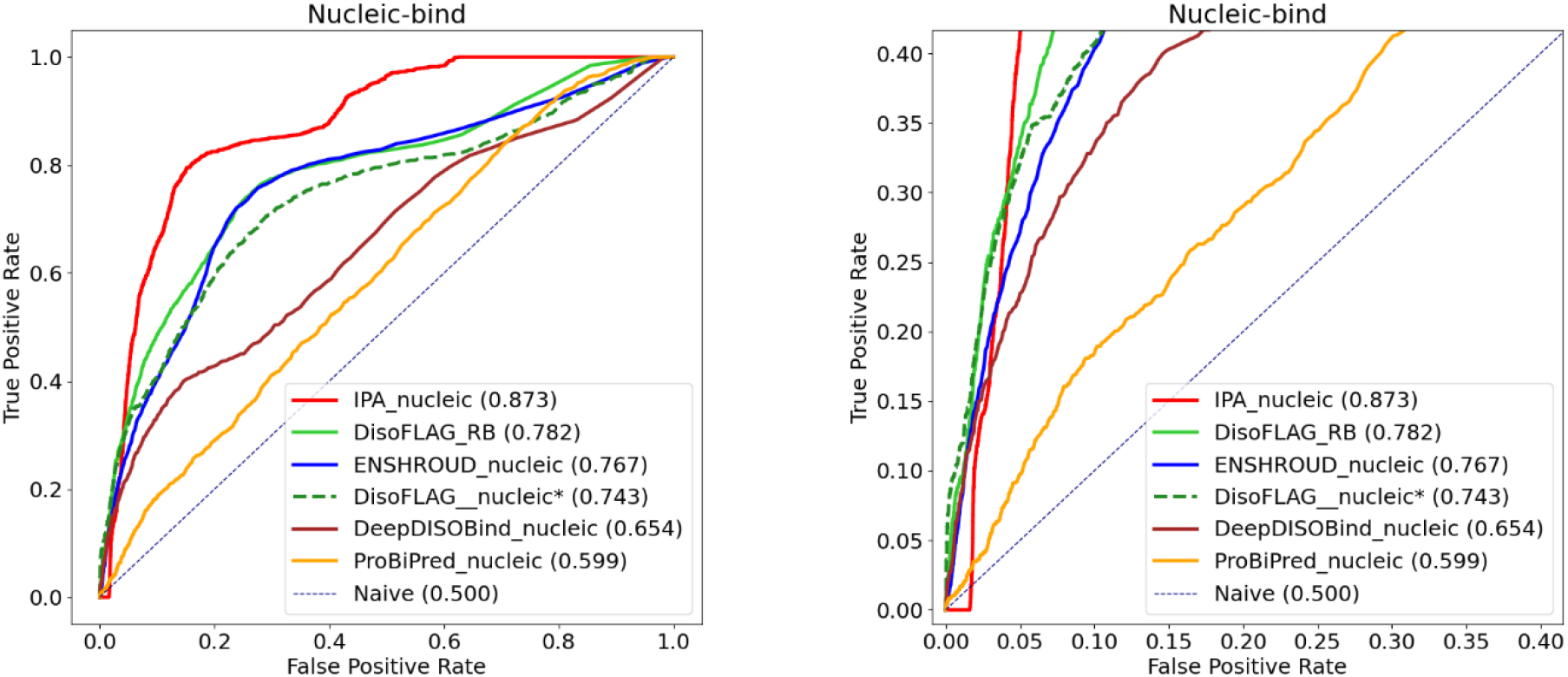
Comparing the full ROC curves (left) of IPA-Nucleic bind predictions to that of DisoFLAG-RB, DisoFLAG-nucleic*, ENSHROUD_nucleic, DeepDISOBind_nucleic, and ProBiPred_nuclic, and a zoom on the lower left corner of these curves (right). (*) The DisoFLAG predictor does not generate predictions for nucleic-binding but generates predictions for RND-binding (RB) and DNA-binding (DB); thus, I averaged the scores of RB and DB as DisoFLAG-nucleic.

Even though IPA-nucleic, with an AUC=0.873, is ahead of other tools and more than 9 points higher than the second tool, DisoFLAF-RB (AUC=0.782), the zoomed-in lower left corner of the ROC curve shows that the IPA-nucleic curve is flat with a TPR=0 for FPR up to 0.0161. That is 1,425 residues annotated as non-nucleic-binding from 13 protein sequences in the Test-24_06 dataset predicted by IPA with a probability of nucleic-binding higher than its prediction of any residue annotated as nucleic-binding, **Table 3**. Even though the test dataset is relatively small, with only 13 binding sites in 11 protein sequences, given the overall IPA-nucleic AUC of 0.873, such a high number of false positives was unanticipated. At first, I contributed this to the UniProt P35637’s 421 residues annotated as non-binding in the IPA-nucleic training data, which represents ∼35% of the 1,183 test nucleic-binding residues, **Figure 2**. However, after investigating these 13 proteins, and in light of our earlier estimation that ∼1% of molecular function sites are annotated, I concluded it is reasonable to assume that many of these 1,425 residues are unannotated nucleic-binding residues. Only six of these 13 proteins in **Table 3** have the UniProt maximum annotation score of 5. Even though UniProt’s maximum annotation score does not imply that proteins are fully annotated, UniProt gave three of these six a molecular function of RNA/DNA binding, which accounts for 456 out of the 599 high-score residues in these six proteins (76%).

**Table 3:**
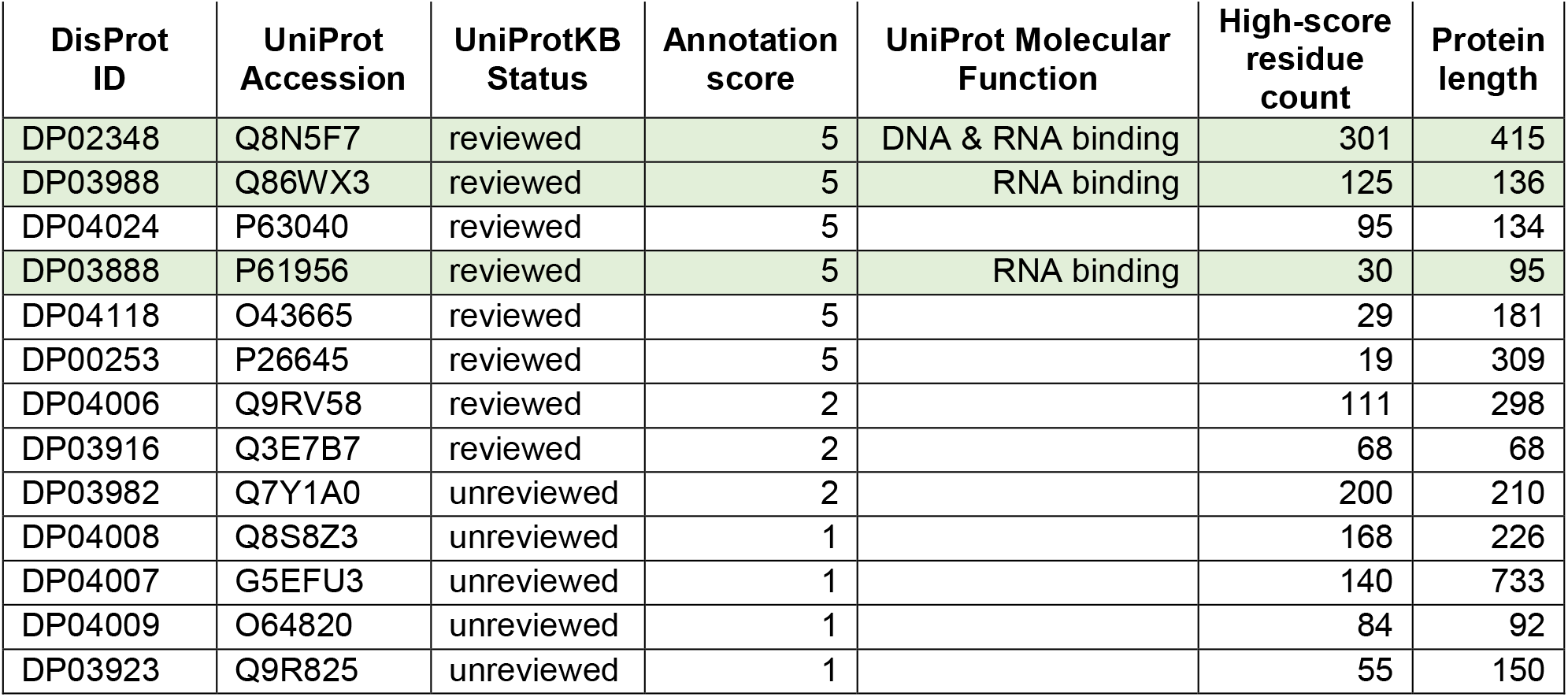
A list of the 13 proteins with high probabilities of IPA nucleic-binding predictions sorted in descending order first by UniProt annotation score and then the number of high-scored residues.

| DisProt ID | UniProt Accession | UniProtKB Status | Annotation score | UniProt Molecular Function | High-score residue count | Protein length |
| --- | --- | --- | --- | --- | --- | --- |
| DP02348 | Q8N5F7 | reviewed | 5 | DNA & RNA binding | 301 | 415 |
| DP03988 | Q86WX3 | reviewed | 5 | RNA binding | 125 | 136 |
| DP04024 | P63040 | reviewed | 5 |  | 95 | 134 |
| DP03888 | P61956 | reviewed | 5 | RNA binding | 30 | 95 |
| DP04118 | O43665 | reviewed | 5 |  | 29 | 181 |
| DP00253 | P26645 | reviewed | 5 |  | 19 | 309 |
| DP04006 | Q9RV58 | reviewed | 2 |  | 111 | 298 |
| DP03916 | Q3E7B7 | reviewed | 2 |  | 68 | 68 |
| DP03982 | Q7Y1A0 | unreviewed | 2 |  | 200 | 210 |
| DP04008 | Q8S8Z3 | unreviewed | 1 |  | 168 | 226 |
| DP04007 | G5EFU3 | unreviewed | 1 |  | 140 | 733 |
| DP04009 | O64820 | unreviewed | 1 |  | 84 | 92 |
| DP03923 | Q9R825 | unreviewed | 1 |  | 55 | 150 |

IPA predictions are fast; using an Intel Core i9, 8/16 cores/threads CPU with 40G DDR4 RAM, and an RTX 3080 GPU machine, it took IPA 4 hours, 9 minutes, and 1 second to process all of the SwissProt (2024_06) 572,619 sequences and generate nucleotide, protein, and all binding predictions. On average, 2,300 sequences per minute or 38.3 sequences per second. When using the CPU to run IPA on the same machine, processing these SwissProt sequences took 11 hours, 45 minutes, and 39 seconds. That is 812 sequences per minute or 13.5 sequences per second.

## 3. Methods

IPA predictors employ a novel window_in/window_out approach. Each predictor utilizes an ensemble of four simple convolutional networks, CNN, models.

### 3.1 CNN model structure

Each CNN consists of three 1D convolutional layers separated by two 1D average pooling layers, followed by two fully connected layers. Thus, the model structure looks like this:

- 1D convolutional layer, with an input size of 510, kernel size CKZ, and depth K.
- ReLU activation function.
- 1D averagePooling layer, with a kernel size equal to APKZ.
- 1D convolutional layer, with a kernel size equal to CKZ and depth (K + 1 * D_inc).
- ReLU activation function.
- 1D averagePooling layer, with a kernel size equal to APKZ.
- 1D convolutional layer, with a kernel size equal to CKZ and depth (K + 2 * D_inc).
- ReLU activation function.
- Fully connected layer 1 (FC1), with an output size FC1Z.
- ReLU activation function.
- Fully connected layer 2 (FC2), with an output size of 10.

Each input sequence is padded at each side by 250 ‘blank’ residues, where a ‘blank’ residue is a hypothetical residue with the values of all its features equal to zero. Then, the first input window starts from the first actual residue at position p=1; we substitute each residue from (p - 250) to (p + 9 + 250) by its K=5 for IPA-nucleic, K=7 features for IPA protein and all, and K=9 features for the IPA-AF2 predictors. The ten outputs of the second fully connected layer, FC2, are associated with input residues positions (p) to (p + 9). Then, we increase p by 10 to generate the second input window and continue repeating these steps until we cover all the sequence residues.

### 3.2 Input Features

Input features are divided into three categories: two affinity features, five DisProt features (**F5** in short), and two alphaFold-2 features. IPA nucleic only utilizes the F5 features, IPA (protein and all) utilizes the affinity and the F5 features, and IPA-AF2 utilizes all nine. Affinity features are derived from P, a 20 × 20 energy predictor matrix [67]. For each position p between 1 and the last in the sequence, one of the affinity features is the sum of the energy in P between the residue at p and the w1=10 residues on each side of p. The second is the sum of this energy between w1 and w2=w1+25, from p + w1 to p + w2 on each p side. The **F5** features are the frequency of each of the 20 standard amino acids in each of the five classes in the DisProt/CAID datasets: IDR, nucleic binding (N_bind), protein binding (P_bind), and ‘Linker’ in the DisProt data and ‘PDB’ annotation in the combined CAID1 and CAID2 sequences, divided by the frequency of that amino acid in the entire dataset, **Supplementary Table S3**. Finally, the two AlphaFold-2 features are those used by the AlphaFold-binding predictor [48] and collected from the AlphaFold-2 generated structures; the alphaFold-2 relative solvent accessibility, RSA, and the AlphaFold-2 prediction accuracy score, pLDDT.

### 3.3 Hyperparameters optimization and the final probability predictions

For all 20 CNN models, four models in each of the five predictors, I used a BCEWithLogitsLoss, thus generating probability outputs. For each model separately, I used a cross-validation grid search to optimize the size of the convolutional kernels (CKZ), the size of the pooling kernels (APKZ), depth increases (D_inc), the output size of the first fully connected layer (FC1Z), and regularization parameters. This results in CKZ between 7 and 15 for all models, APKZs are 7 or 15, D_incs are 2 or 10, and FC1Zs are between 100 and 150. A complete list of values for CKZ, APKZ, D_inc, and FC1Z values for all 20 CNN models is available in **Supplementary Table S4**.

For each predictor, first, I applied a reversed Bayes Rule to the probability output to factor out the priors of its training data; thus, the scores for all four outputs were aligned, and then I averaged these four scores to produce the predictor probability output.

## Discussion

This paper presents a new platform for the probability annotation of intrinsically disordered proteins (IPA). The IPA platform includes tools designed to predict nucleotide, protein, and all binding sites within intrinsically disordered regions (IDRs) using only amino acid sequences. A separate tool, IPA-AF2, also leverages AlphaFold-2 structural models to predict protein binding sites and linker regions. To evaluate the performance of IPA, I constructed a test dataset that is completely homology-free to IPA training data. Results demonstrate that despite relying solely on amino acid sequences and achieving high computational efficiency, IPA favorably matched leading tools in predicting protein and all IDR binding sites, and it significantly outperforms in identifying nucleic binding sites and linker regions.

Most importantly, I introduced a new output paradigm form for computational predictions of features. Currently, the output of tools predicting features is expressed as a score, where higher values indicate a greater probability. Some tools rely on heuristics to generate these scores [48][37], while others heuristically redistribute them to fit some arbitrarily selected distribution [31][34]. Although most are in the range between zero to one, these scores are not actual probabilities even when logistic regression is used, as the priors of the training data do not reflect the unknown priors of the general population. As a result, these scores are not comparable across different tools. They are also meaningless independently and can only be interpreted by comparing them to others of the same tool. As an alternative, I am suggesting the application of a reverse Bayes Rule to the output of logistic regression models to factor out the priors in the training data. As a result, the final prediction scores will reflect probabilities, assuming 50% positive in the general population. Thus, these scores are individually interpretable, and if adopted by other tools, the output of these tools can be compared and integrated.

## Notes

### Competing Interest Statement

The authors have declared no competing interest.

### Summary of Updates

In the original script, the first two figures were named Figure 1. A small number of other typos

