## Supplement Figure S1, and tables S1, S2, S3, & S4 for "Probabilistic Annotations of Protein Sequences for Intrinsically Disordered Features"

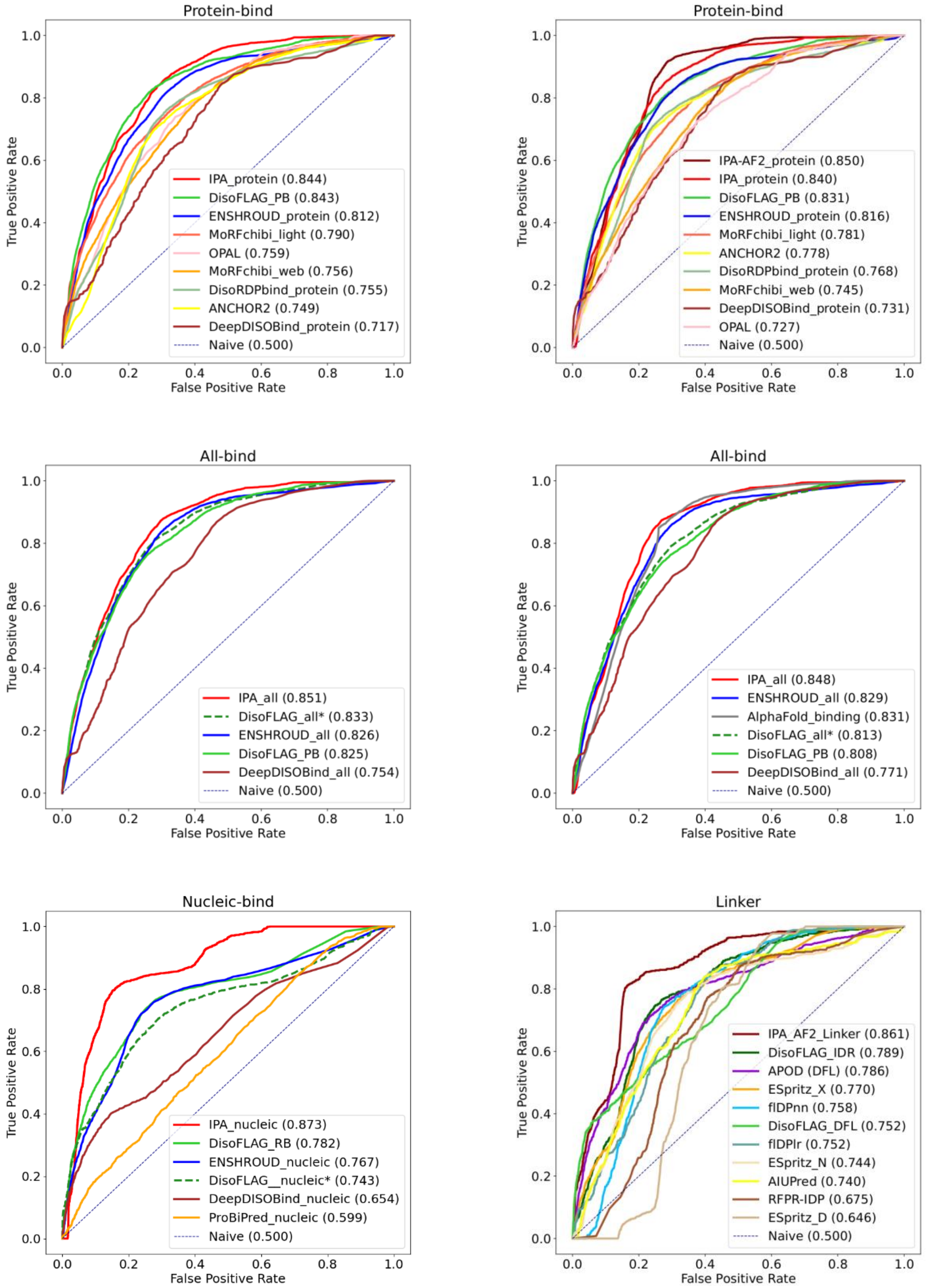
Protein-bind
True Positive Rate
False Positive Rate
IPA_protein (0.844)
DisoFLAG_PB (0.843)
ENSHROUD_protein (0.812)
MoRFchibi_light (0.790)
OPAL (0.759)
MoRFchibi_web (0.756)
DisoRDPbind_protein (0.755)
ANCHOR2 (0.749)
DeepDISOBind_protein (0.717)
Naive (0.500)
Protein-bind
True Positive Rate
False Positive Rate
IPA-AF2_protein (0.850)
IPA_protein (0.840)
DisoFLAG_PB (0.831)
ENSHROUD_protein (0.816)
MoRFchibi_light (0.781)
ANCHOR2 (0.778)
DisoRDPbind_protein (0.768)
MoRFchibi_web (0.745)
DeepDISOBind_protein (0.731)
OPAL (0.727)
Naive (0.500)
All-bind
True Positive Rate
False Positive Rate
IPA_all (0.851)
DisoFLAG_all* (0.833)
ENSHROUD_all (0.826)
DisoFLAG_PB (0.825)
DeepDISOBind_all (0.754)
Naive (0.500)
All-bind
True Positive Rate
False Positive Rate
IPA_all (0.848)
ENSHROUD_all (0.829)
AlphaFold_binding (0.831)
DisoFLAG_all* (0.813)
DisoFLAG_PB (0.808)
DeepDISOBind_all (0.771)
Naive (0.500)
Nucleic-bind
True Positive Rate
False Positive Rate
IPA_nucleic (0.873)
DisoFLAG_RB (0.782)
ENSHROUD_nucleic (0.767)
DisoFLAG__nucleic* (0.743)
DeepDISOBind_nucleic (0.654)
ProBiPred_nucleic (0.599)
Naive (0.500)
Linker
True Positive Rate
False Positive Rate
IPA_AF2_Linker (0.861)
DisoFLAG_IDR (0.789)
APOD (DFL) (0.786)
ESpritz_X (0.770)
flDPnn (0.758)
DisoFLAG_DFL (0.752)
flDPlr (0.752)
ESpritz_N (0.744)
AIUPred (0.740)
RFPR-IDP (0.675)
ESpritz_D (0.646)
Naive (0.500)

**Figure S1:** Compared to other tools, the ROC curves for the IPA protein, nucleic, and all binding predictions and IPA-AF2 protein-binding and Linker regions predictions. The three ROC curves in the left column are generated against the **Test-24_06** dataset and those in the right column against the **Test-24_06-AF2**. (*) The DisoFLAG predictor does not generate predictions for nucleic-binding and all-binding but generates predictions for RND-binding (RB), DNA-binding (DB), and protein-binding (PB); thus, I averaged the scores of RB and DB as DisoFLAG_nucleic, and then averaged DisoFLAG_nucleic with PB as DisoFLAG_all.

**Table S1:** Steps used to assemble the Test-24_06 dataset. Test-24_06-AF2 is a subset of the TEST-24_06 sequences with structures generated by AlphaFold-2 in the AlphaFold database.

| | | **Total sequences** | **Annotated sequences** | **Target sites** | **Non-target residues** | **Target residues** | **Masked-out residues** |
|---|---|---|---|---|---|---|---|
| **DSET1** = DP24_06 - DP23_12 | **P_bind** | 185 | 25 | 28 | 96,714 | 2,910 | 0 |
| | **N_bind** | 185 | 5 | 5 | 99,212 | 412 | 0 |
| | **LIP** | 185 | 28 | 31 | 96,470 | 3,154 | 0 |
| | **Linker** | 185 | 14 | 19 | 98,981 | 643 | 0 |
| **DSET2** = DSET1 after excluding homologs with > 30% identity using CD-Hit | **P_bind** | 164 | 19 | 22 | 80,966 | 2,224 | 8,505 |
| | **N_bind** | 164 | 5 | 5 | 88,324 | 412 | 2,959 |
| | **LIP** | 164 | 22 | 25 | 80,722 | 2,468 | 8,505 |
| | **Linker** | 164 | 12 | 16 | 86,543 | 559 | 4,593 |
| **DSET3** = Including MobiDB annotations into DSET2 | **P_bind** | 164 | 26 | 33 | 68,778 | 2,515 | 20,402 |
| | **N_bind** | 164 | 5 | 5 | 88,324 | 412 | 2,959 |
| | **LIP** | 164 | 40 | 51 | 67,264 | 4,147 | 20,284 |
| | **Linker** | 164 | 12 | 16 | 86,543 | 559 | 4,593 |
| **DSET4** = DSET3 after processing with HAM | **P_bind** | 164 | 26 | 33 | 68,719 | 2,515 | 20,461 |
| | **N_bind** | 164 | 5 | 5 | 88,279 | 412 | 3,004 |
| | **LIP** | 164 | 40 | 51 | 67,060 | 4,147 | 20,488 |
| | **Linker** | 164 | 12 | 16 | 86,521 | 559 | 4,615 |
| **Test-24_06** = DSET4 + DP23_12 sequences with new annotations | **P_bind** | 184 | 42 | 55 | 68,719 | 3,873 | 28,742 |
| | **N_bind** | 184 | 11 | 13 | 88,279 | 1,183 | 11,872 |
| | **LIP** | 184 | 48 | 65 | 68,445 | 4,696 | 28,193 |
| | **Linker** | 184 | 14 | 18 | 86,521 | 619 | 14,194 |
| **Test-24_06-AF2 (limited to AlphaFold2 structures)** | **P_bind** | 147 | 35 | 47 | 58,985 | 3,436 | 13,123 |
| | **LIP** | 147 | 40 | 55 | 56,567 | 4,220 | 11,757 |
| | **Linker** | 144 | 14 | 18 | 67,519 | 619 | 584 |

**Table S2:** IPA-AF2_Linker predictor outperformed other Linker predictors (in bold) and IDR predictors in AUC values against the TEST-24_06-AF2 dataset.

| | **AUC** |
|---|---|
| **IPA_AF2_Linker** | **0.861** |
| DisoFLAG_IDR | 0.789 |
| **APOD** | **0.786** |
| ESpritz_X | 0.770 |
| flDPnn | 0.758 |
| **DisoFLAG_DFL** | **0.752** |
| flDPlr | 0.752 |
| ESpritz_N | 0.744 |
| AIUPred | 0.740 |
| RFPR-IDP | 0.675 |
| ESpritz_D | 0.646 |

**Table S3:** The **F5** features for each of the 20 standard amino acids. For each amino acid, these features are the frequency of that amino acid in each of the five classes: IDR, nucleic binding (N_bind), protein binding (P_bind), and 'Linker' in the DisProt 2023_12 data and 'PDB' annotation in the combined CAID1 and CAID2 sequences, divided by the frequency of that amino acid in the entire dataset and then multiplied by 100.

| Amino Acid | IDR | Linker | N_bind | P_bind | PDB |
|---|---|---|---|---|---|
| A | 108.776 | 105.711 | 96.964 | 103.172 | 93.301 |
| C | 51.852 | 39.502 | 51.072 | 59.233 | 117.768 |
| D | 112.017 | 109.835 | 97.596 | 113.697 | 101.451 |
| E | 124.133 | 114.155 | 104.308 | 118.074 | 96.901 |
| F | 69.997 | 59.429 | 73.528 | 74.956 | 115.953 |
| G | 116.862 | 121.453 | 111.039 | 111.067 | 94.527 |
| H | 83.513 | 79.084 | 91.107 | 80.180 | 98.521 |
| I | 66.664 | 67.150 | 81.632 | 73.633 | 121.547 |
| K | 119.192 | 102.862 | 158.014 | 100.009 | 101.104 |
| L | 71.549 | 65.828 | 66.448 | 83.231 | 109.540 |
| M | 82.179 | 92.349 | 77.203 | 84.958 | 100.098 |
| N | 97.746 | 117.902 | 114.751 | 100.493 | 103.319 |
| P | 133.391 | 147.432 | 110.458 | 126.119 | 78.845 |
| Q | 112.595 | 123.824 | 113.074 | 114.435 | 97.290 |
| R | 89.594 | 91.773 | 127.314 | 100.377 | 102.433 |
| S | 117.347 | 138.093 | 116.409 | 122.400 | 82.040 |
| T | 106.789 | 108.159 | 96.611 | 104.118 | 92.888 |
| V | 81.305 | 72.932 | 73.531 | 81.647 | 106.683 |
| W | 62.060 | 42.601 | 53.834 | 65.289 | 121.847 |
| Y | 69.566 | 50.065 | 85.432 | 72.148 | 119.810 |

**Table S4:** The complete list of values for convolutional kernel sizes (CKZ), average pooling kernel sizes (APKZ), depth increase (D_inc), and the output size of the first fully connected layer (FC1Z) values for all 20 CNN models

| Predictor | Model CV num | CKZ | APKZ | D_inc | FC1Z |
|---|---|---|---|---|---|
| IPA-Protein | 1 | 10 | 15 | 4 | 100 |
| | 2 | 15 | 15 | 6 | 100 |
| | 3 | 7 | 15 | 4 | 150 |
| | 4 | 15 | 15 | 4 | 100 |
| IPA-Nucleic | 1 | 10 | 15 | 8 | 150 |
| | 2 | 7 | 10 | 4 | 100 |
| | 3 | 7 | 15 | 6 | 100 |
| | 4 | 15 | 7 | 4 | 100 |
| IPA-All | 1 | 10 | 10 | 10 | 130 |
| | 2 | 10 | 10 | 10 | 130 |
| | 3 | 10 | 10 | 8 | 130 |
| | 4 | 10 | 10 | 8 | 110 |
| IPA-AF2-Protein | 1 | 7 | 10 | 6 | 100 |
| | 2 | 10 | 15 | 4 | 150 |
| | 3 | 7 | 15 | 4 | 100 |
| | 4 | 15 | 10 | 4 | 150 |
| IPA-AF2-Linker | 1 | 10 | 10 | 2 | 100 |
| | 2 | 10 | 15 | 4 | 150 |
| | 3 | 7 | 7 | 4 | 150 |
| | 4 | 15 | 10 | 4 | 100 |
